# Proliferation/Differentiation in intestinal organoids as a balance of ligand-modulated EGFR traffic

**DOI:** 10.64898/2026.03.30.715070

**Authors:** MO Caracci, S Seidler, LM Muñoz-Nava, B Soetje, K Michel, PIH Bastiaens

**Author notes:** Deceased (PIH Bastiaens).

## Abstract

Epidermal Growth factor (EGF) signaling is associated with (oncogenic) proliferation. Conversely, EGF-family ligands are able to trigger a differentiation program in cultured cells, an effect attributed to ligand affinity and EGFR phosphorylation. How EGF/EGFR driven proliferation-differentiation dynamics underlie tissue self-renewal has not been addressed. We show that culturing mouse small intestinal organoids (mSIOs) without EGF enhanced EGFR expression and base phosphorylation while maintaining a balanced development of proliferative crypts and differentiated villi. Addition of EGF or EREG triggers receptor endocytosis, reducing cell-surface and expression levels. While EGF promoted crypt proliferation, EREG promoted both proliferation and villus differentiation compared to untreated controls. Removal or re-introduction of EGF or EREG proved sufficient to induce development comparable to constant presence of ligands over 96h. Sub-saturating concentrations of EGF led to increased villus differentiation, resembling EREG treatments, suggesting that control over EGFR endocytic cycle ultimately regulates the balance of proliferation and differentiation in mSIOs

**Summary:** Expression and signaling competency at the plasma membrane of EGFR drives crypt proliferation vs villus differentiation by medium ligand-composition, aiding mouse intestinal organoids self-renewal and regeneration.

## Introduction

Intestine morphology is characterized by differentiated enterocytes in the villus regions extending into the chyme for nutrient uptake, and a proliferative stem cell niche protected at bottom of crypts. Here, EGF/EGFR signaling is essential for maintance proliferation and self-renewal of the intestinal stem cells. Aberrant activity of EGF downstream signaling has been consistently associated with oncogenic transformations. EGF is a fundamental growth factor for many in vitro organoid culture systems (van der Vaart et al., 2021; Mahe et al., 2013; Wang et al., 2025; Hu et al., 2018). In mouse small intestinal organoids (mSIOs), Paneth cells secrete paracrine EGF within the stem cell niche in crypts (Sato et al., 2011a). Nevertheless, additional growth factors, including EGF, are added to the culture medium, mimicking the complexity of stromal compartment surrounding the intestinal epithelia (Sato et al., 2009).

EGFR is mainly expressed in the intestinal crypts (Yang et al., 2017) and inhibition of tyrosine kinase activity in mature organoids led to cell cycle arrest and quiescence of LGR5+ stem cells (Basak et al., 2017). EGFR inhibition also enhanced differentiation towards entero-endocrine cells, suggesting that perturbations to EGF/EGFR activation during distinct developmental stages could aid differentiation in mSIOs. Other EGF-family ligands such as NRG1, EREG and HB-EGF can replace EGF in mSIOs culture promoting organoid development (Lemmetyinen et al., 2023; Jardé et al., 2020; Chen et al., 2012), notably epithelial KO of NRG1 does not affect differentiation of secretory linage cells(Jardé et al., 2020). While NRG1 does not bind EGFR, EREG binds both EGFR and ERBB4 and has been used in human intestinal organoids to enhance cellular differentiation and crypt development (Childs et al., 2023).

Under basal medium conditions, mSIOs show both crypt proliferation and villus differentiation in two distinct, spatially separated compartments. In contrast, human intestinal organoids feature less cellular and structural diversity. Several remarkable efforts have been taken to enhance human organoid complexity by modulating cell signaling, including EGF inhibitors and EGF-family ligands. Nevertheless, the spatio-temporal responses of dynamic intracellular signaling networks to extracellular ligands have been largely overlooked. (Yang et al., 2025; Fujii et al., 2018; Sato et al., 2011b; He et al., 2022).

It is well stablished that EGFR signaling plays an integral role in partially opposing phenotypic outcomes, including migration, apoptosis, proliferation and differentiation and is generally acknowledged that these responses are a result of different affinities among EGF-family ligands and their capacity to induce receptor phosphorylation (Abud et al., 2021; Jendrisek et al., 2026). For instance, high affinity ligands such as EGF, HB-EGF and TGFa, all produced in the intestinal epithelia, are often associated to cell proliferation. In other hand, low affinity ligands such as EREG and AREG, which are expressed by stromal cells in the intestine, are associated to cell differentiation and migration (Jendrisek et al., 2026; Lemmetyinen et al., 2023). Interestingly, EGF triggered responses are dependent on ligand concentration, where high, receptor saturating concentrations promote cellular proliferation while low, sub-saturating concentrations promote migratory behavior in cultured cells (Brüggemann et al., 2021; Joshi et al., 2023). The fact the EGF-EGFR signaling is able to trigger multiple phenotypical responses underscores the importance of this ligand-receptor pair in development. However, these properties have mainly been demonstrated in immortalized cell lines and this decision-taking has not yet been explored in complex tissues under homeorhetic conditions.

Therefore, we studied mSIOs in their early developmental stages during self-renewal and regeneration of seeded single crypts and their subsequent maturation. For this, we cultured mSIOs with either EGF or EREG to examine the differential effect of ligand availability on sustaining organoid crypt and villus development in aiding homeorhetic regeneration. We find that maintenance of plasma membrane (PM) EGFR localization drives villus development regardless of ligand affinity.

## Results/Discussion

### EGF promotes crypt proliferation and EREG promotes villus differentiation

In order to understand how the balance within crypt proliferation and villus differentiation is triggered by distinct EGF-family growth factors we stained organoids with CD44, a bonafide crypt marker (Sumigray et al., 2018), and enterocyte marker AldoB to identify the differentiated villus region (Serra et al., 2019). Images were acquired by CLSM and maximum intensity projections from multiple confocal stacks were masked and quantified for area, as surrogates for crypt and villus growth triggered by our treatments (Figure 1A and B). Intestinal organoid crypts were grown for 96h in control NR medium (Noggin, R-SPO1) or NR medium supplemented with either EGF (50 ng/mL), EREG (50 ng/mL). EGF and EREG as high and low affinity ligands of EGFR respectively, are known to induce receptor phosphorylation and endocytosis (Freed et al., 2017).

**Figure 1.**
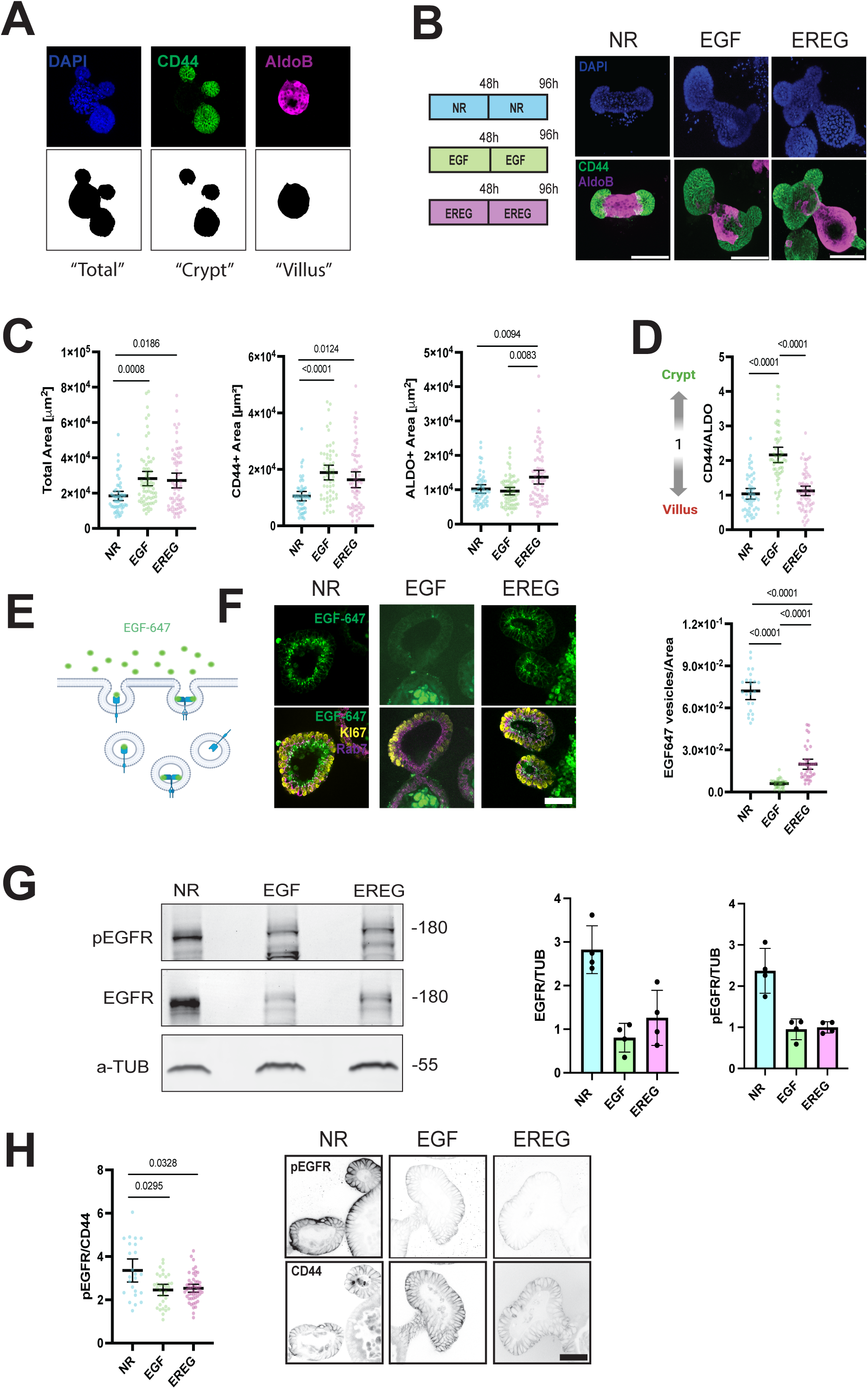
EGFR surface depletion by EGF promotes Crypt proliferation while EREG promotes both proliferation and differentiation. (A) Maximum intensity projections (top) were binarized (bottom) to quantify the area of DAPI (nuclei, blue), CD44 (Green) and AldolaseB (Magenta) used to quantify “Total”, “Crypt” and “Villus” area respectively (B-H) Organoids were grown, with NR medium or supplemented with either EGF (50ng/mL) or EREG (50ng/mL) for 96h with a complete medium change at 48h. (B) Organoids were immuno-stained with CD44 (Green) AldoB (Magenta) and DAPI (Blue). Scale Bar 100 μm. (C) Quantification of the areas as shown in (A). Scatter plots with mean±95%CI. N=3; n=57-67 organoids/condition; significance was calculated by Kruskal-Wallis with Dunn’s multiple comparisons test. p-values <0.05 are shown (D) Relative ratio of CD44 to ALDO areas from (C). Scatter plot with mean±95%CI; significance was calculated by Kruskal-Wallis with Dunn’s multiple comparisons test. p-values <0.05 are shown. Values above 1 suggests organoid development tends towards crypt proliferation while values below 1 tend to villus differentiation (E) Alexa Fluor-647 labelled EGF (50 ng/mL) is added to organoids after 96h of growth, after 30 min of incubation organoids were fixed and stained. Alexa Fluor-647 positive endosomes indicate the level of surface EGFR and consecutive endocytosis upon ligand binding. (F) (Left) Representative micrographs of organoids after immunofluorescence staining against Ki67 (Yellow) and Rab7 (Magenta). EGF-647 internalized through EGFR endocytosis is labelled in (Green). Scale bar 20μm. (Right) EGF-647 vesicles per area as scatter plots mean±95%CI; N=3, n=28-37 crypts/condition; significance was calculated by Kruskal-Wallis with Dunn’s multiple comparisons test. p-values <0.05 are shown. (G) (Left) Immunoblot of pEGFR, EGFR and a-TUB with approximate MW in kDa. (Right) Bar Graphs of mean±SD of pEGFR or EGFR relative to a-TUB. N=4. (H) (Left) Ratio of pEGFR to CD44 Scatter plots of mean±95%CI. N=3, n=32-60 crypts/condition; significance was calculated by Kruskal-Wallis with Dunn’s multiple comparisons test. p-values <0.05 are shown. (Right) Representative micrographs of organoids after immunofluorescence staining against pEGFR (top) and CD44 (bottom). Scale bar 20μm.

EGF and EREG addition significantly enhanced Organoid growth compared to NR medium, however, Crypt Area was differentially enhanced by EGF, while Crypt and Villus area was enhanced by EREG (Figure 1C) The relative ratio of Crypt-to-Villus (CD44/ALDO) showed that EGF strongly favored the Crypt proliferation over Villus differentiation, while NR medium and EREG maintained balanced development of Crypt and Villus, despite the overall larger size of EREG treated organoids relative to NR treatment (Figure 1D).

Crypt number was also greatly enhanced by EGF, while EREG only had small effect in increasing the number of crypts relative to NR organoids (Figure S1A). Crypt establishment is a direct result of apical constriction by Paneth cells(Langlands et al., 2016). Paneth cells are found in the crypt, interspaced between Intestinal Stem cells (ISCs) and their abundance is often correlated to ISCs numbers (Tallapragada et al., 2021). Immuno-fluorescence staining showed that neither EGF or EREG enhanced the number of Paneth cell, regardless of the effect observed in crypt area (Figure S1B).

Results suggests that both EGF and EREG are capable of inducing organoid development through distinctive regulation of the proliferation/differentiation balance with EGF promoting CD44+ crypts enlargement over differentiation into ALDO+ villus region. On the other hand, EREG promotes growth of both crypt and villus regions, thus suggesting that EGF-family ligands promote mSIOs development through distinctive downstream mechanisms.

### EGF and EREG mediated endocytosis reduces EGFR PM localization and expression

EGF and EREG are known to bind, induce endocytosis and phosphorylation of EGFR. EREG is also a ligand of ERBB4, however expression of this receptor is not detected in small intestinal epithelia or stroma (Lemmetyinen et al., 2023). Saturating EGF concentrations are commonly used in Organoid culture (Shankaran et al., 2021)at which most of the PM receptor is internalized and undergo lysosomal degradation as demonstrated in cultured immortalized cells (Brüggemann et al., 2021). In contrast, EREG binding to EGFR promotes recycling back to the plasma membrane (Roepstorff et al., 2009). In order to quantify end-point surface levels of EGFR, organoids were incubated for 30 min with 50 ng/mL of EGF-647 before fixation and staining (Figure 1E). We observe that most of the internalized EGF-647 is located in the crypt regions with few vesicles being observable in the villus regions, this is consistent with the literature stating that EGFR is highly expressed in cycling cells within the intestinal crypt (Yang et al., 2017). EGF-647 endocytosis was observed in organoids grown in NR or EREG medium, organoids grown with unlabeled EGF for 96h showed very low endocytosis suggesting reduced levels of PM EGFR (Figure 1F). Absence of ligands allowed for accumulation of surface EGFR throughout the 96h developmental window and EREG mediumted recycling back to the plasma membrane maintained high surface levels.

EGF-647 endocytosis showed co-localization with late endosome marker Rab7 consistent with the use of saturating amounts of ligand (Figure 1F; Figure S2A and B). DLS (Digital Lightsheet) imaging of EGF-647 endocytosis shows an initial association to the crypts basal membrane followed by the entrance of vesicles travelling adjacent to the inner membranes towards the apical side (Figure S2C and Video1). While basal to apical trafficking is clearly observed we cannot discard the possibility of apical endocytosis of EGF-647 from the lumen, as cell extrusions within the crypts are observable in the time frame of our experiments which could allow EGF-647 to seep into the luminal compartment. Transcytosis of EGF from basal to apical has been proposed in MDCK epithelial layers (Brändli et al., 1991), however little is known of the mechanisms involved As a PM receptor, EGFR phosphorylation is highly dependent on its interaction with ligands in the extracellular medium, nevertheless gene amplification enhancing expression levels or point mutations affecting inhibitory mechanisms can lead to EGFR phosphorylation in absence of ligand (Stanoev et al., 2018; Arteaga and Engelman, 2014). Immunoblotting of both total and phosphorylated EGFR Y1068 showed that EGF and EREG grown organoids had a significant reduction in EGFR protein expression compared to NR grown organoids (Figure 1G). Immuno-fluorescence staining pEGFR Y1068 and CD44 after treatments, showing an increase in phosphorylation staining only in crypts growth without EGF or EREG while organoid grown with either ligand showed reduced staining (Figure 1H).

EGFR endocytic cycle is fundamental to regulate receptor phosphorylation, allowing for degradation in late endosomes or the interaction with protein tyrosine phosphatases (PTPs) within endomembrane compartments (Baumdick et al., 2015; Joshi et al., 2023; Stanoev et al., 2018). In turn, NR grown organoids show an accumulation of pEGFR likely due to endogenously expressed EGF by Paneth cells and amplification by autocatalytic activation of PM bound receptors. It is important to consider that medium changes every 48h imply 2 “pulses” of fresh EGF or EREG in a 96h developmental window, in between pulses, biosynthetic and endocytic mechanisms allow for the replenishment of surface EGFR (Scharaw et al., 2016).Surprisingly while EREG treatment did allow for significantly higher EGFR PM levels, total levels were greatly diminished, despite this ligand promoting receptor recycling over late endosome trafficking (Figure 1G and H).

### Temporal modulation of EGF-family ligands during development alters villus differentiation

In vitro cell culture often promotes proliferating conditions which ultimately allow for biomass expansion required for biochemical assays, and organoid culture is no exception. In the conception of ENR medium, addition of EGF and R-SPO1 were sufficient to induce organoid growth from single crypts and the addition of Noggin proved instrumental for organoid passaging over time (Sato et al., 2009). While lower concentrations of EGF (e.g 10 ng/mL) were also tested in this seminal work with similar results, little consideration was given to organoid developmental paths under evolving availability of EGF in regards to ligand concentration or duration of stimuli.

To test the temporal requirements of EGF/EGFR signaling in organoid development, organoids were grown for 96h with either EGF or EREG. However a complete medium exchange with NR medium (No EGF or EREG) was done after 1h or after 48h of incubation (Figure 2A). Developing from single crypts, organoids activate a regeneration program that initially drives proliferation into a fetal-like state where differentiation and stem cells markers are mostly absent (Sprangers et al., 2021), budding within the organoid usually appears at the 48h mark where mature crypts will develop (Sato et al., 2009) (Figure S3A). After 48h in culture, most organoids feature a round morphology that is entirely CD44 positive, consistent with an undifferentiated proliferating organoid (Serra et al., 2019). About 25% of them showed AldoB staining, suggesting initial establishment of a villus region. The percentage of organoids showing AldoB staining increases with EREG treated crypts, as shown for 96h organoids grown with EREG. Overall, total area of organoids was not changed in any of the treatments regardless of presence or absence of AldoB staining (Figure S3B-E). EGF-647 endocytosis was also clearly observed in NR and EREG grown organoids but not in EGF grown organoids as shown for 96h organoids. Endocytosis was possible in organoids displaying either crypt or round morphology, often confined to proliferating cells as noted by Ki67 staining (Figure S3F).

**Figure 2.**
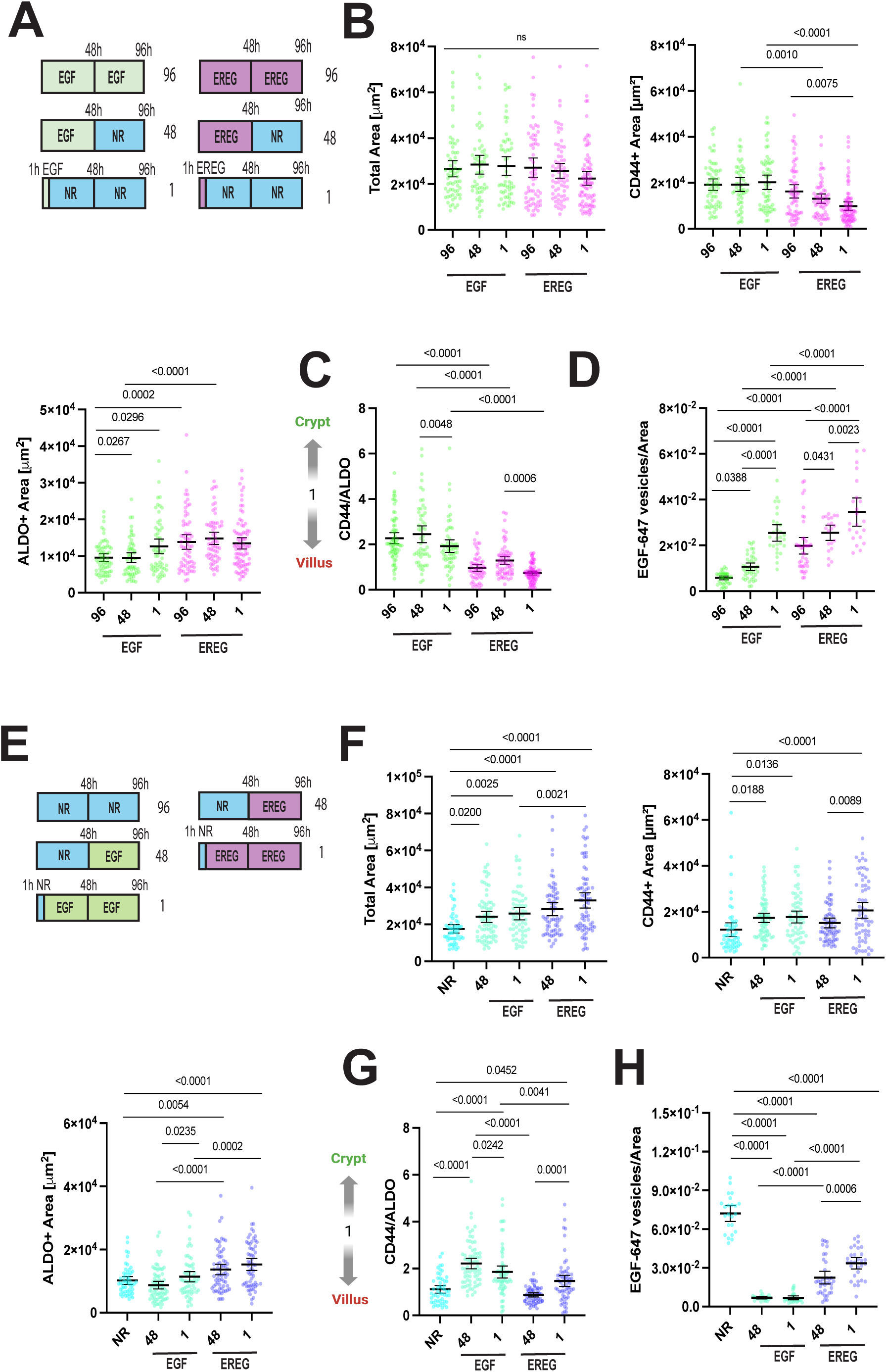
Removing EGF or EREG after crypt seeding does not modify developmental trayectories. (A-D) Organoids were grown, with NR medium supplemented with either EGF (50ng/mL) or EREG (50ng/mL) for 48h or 1h, then complete medium was replaced with NR medium until fixation at 96h. (B) Scatter plots showing mean±95%CI, N=3, n=57-79 organoids/condition of masked Total, CD44+ and ALDO+ areas. Significance was calculated by Two-way ANOVA and Sidak’s multiple comparisons test multiple comparisons test. p-values <0.05 are shown. (C) Relative ratio of CD44 to ALDO areas from (B). Scatter plot mean±95%CI of relative ratio within CD44/ALDO areas. Values above 1 suggests organoid development tends towards crypt proliferation while values below 1 tend to villus differentiation. Significance was calculated by Two-way ANOVA and Sidak’s multiple comparisons test. p-values <0.05 are shown. (D) EGF-647 vesicles per area as scatter plots showing mean±95%CI; N=3, n=28-41 crypts/condition; significance was calculated by Two-way ANOVA and Sidak’s multiple comparisons test multiple comparisons test. p-values <0.05 are shown. (E-H) Organoids were grown for 1h and 48h with NR, then complete medium was replaced with NR medium supplemented with either EGF (50ng/mL) or EREG (50ng/mL). (F) Scatter plots showing mean±95%CI, N=3, n=57-73 organoids/condition of masked Total, CD44+ and ALDO+ areas. Significance was calculated by Two-way ANOVA and Sidak’s multiple comparisons test multiple comparisons test. p-values <0.05 are shown, ns non-significant. (G) Relative ratio of CD44 to ALDO areas from (B). Scatter plot mean±95%CI of relative ratio within CD44/ALDO areas. Values above 1 suggests organoid development tends towards crypt proliferation while values below 1 tend to villus differentiation. Significance was calculated by Two-way ANOVA and Sidak’s multiple comparisons test multiple comparisons test. p-values <0.05 are shown. (H) EGF-647 vesicles per area as scatter plots showing mean±95%CI; N=3, n=20-38 crypts/condition; significance was calculated by Two-way ANOVA and Sidak’s multiple comparisons test multiple comparisons test. p-values <0.05 are shown.

Regardless of EGF or EREG removal time-frame, organoids showed comparable total area after 96h in culture. Moreover, 1h and 48h of EGF containing medium was sufficient to induce crypt growth comparable to 96h with constant EGF. EREG induced villus differentiation showed a similar trend as EGF with crypt development. Surprisingly, 1h EGF treatment not only induced crypt growth, but also enhanced villus differentiation to levels similar to the ones observed in EREG treatment (Figure2B and C). Removal of EGF or EREG did not affect crypt number, however, removal of EGF increased Paneth cell number compared to 96h control (FigureS1C and D). Alternatively, organoids were treated for 96h and 1h with 50ng/mL of NRG1 (Figure S1I), An EGF-family ligand know to bind ERBB3 and ERBB4 but not EGFR. NRG1 was able to induce organoid growth as previously shown (Jardé et al., 2020; Lemmetyinen et al., 2023). Notably, 1h treatment with NRG1 and subsequent withdrawal failed to induce organoid growth to 96h levels, with a significant growth-increase relative to NR-grown organoids (Figure S1J and K). Therefore EGFR is likely to induce robust signaling responses under EGF and EREG treatments.

96h and 48h incubation with EGF led to a near complete depletion of surface EGFR, as very low EGF-647 endocytosis was detected., In contrast, 1h incubation with EGF exhibited high levels of endocytosis suggesting high levels of surface EGFR, similar to EREG treatments which increased over EREG-starvation times (Figure 2D). EGFR phosphorylation was enhanced by EGF or EREG removal compared to the 96h treatments further highlighting that phosphorylation of EGFR requires PM localization (Figure S1E).

In a complementary experiment, single crypts were seeded in NR medium and then ligands were re-introduced after 48h or 1h of growth factor starvation. (Figure 2E). EGF and EREG-containing medium increased organoids total area and crypt area. In contrast to early EREG treatment, addition of EREG after growth factor starvation enhanced crypt growth similar to EGF and at the same time increased villus growth as previously established (Figure 2F-G). Likewise, there was a marked increase in the number of organoids featuring multiple crypts, although only EGF treatment enhanced the number of Paneth cells (Figure S1F and G).

As previously shown, EGF and EREG reintroduction into the culture medium greatly reduced endocytosis of labelled EGF-647 with EREG treated organoids retaining some endocytic capacity consistent with EGFR recycling towards the plasma membrane (Figure 2H). EGF or EREG removal from the culture medium allowed for increased phosphorylation of EGFR, likewise, re- introducing EGF or EREG at 48h after seeding significantly reduced phosphorylation, while brief 1h starvation did not change phosphorylation levels compared to the NR condition (Figure S1H).

Altogether, these results highlight the remarkable robustness of EGFR signaling in response to high affinity and low affinity ligands, effect that could not be replicated by NRG1 treatment, which does not bind EGFR. EGF or EREG treatment at any point tested was able to reduce EGFR surface levels compared to NR and lead to similar total area growth, with 1h treatments upon seeding being sufficient to induce crypt and villus development respectively.

### Sub-saturating concentrations of EGF enhance both crypt proliferation and villus differentiation

Given that organoids can develop in EGF-free medium, albeit at smaller size of crypt and villus; and that EREG, a low affinity ligand of EGFR, induces organoid development while maintaining high levels of surface EGFR, we tested if lower, non-saturating concentrations of EGF could sustain organoid growth. Single crypts were cultured for 96h with EGF concentrations decreasing by one order of magnitude of starting at 50 ng/mL to 0.05 ng/mL (Figure 3A). NR treatment (0 ng/mL of EGF) showed consistent development compared to 50 ng/mL as shown in previous experiments, however, concentrations in between showed remarkable differences in development.

**Figure 3.**
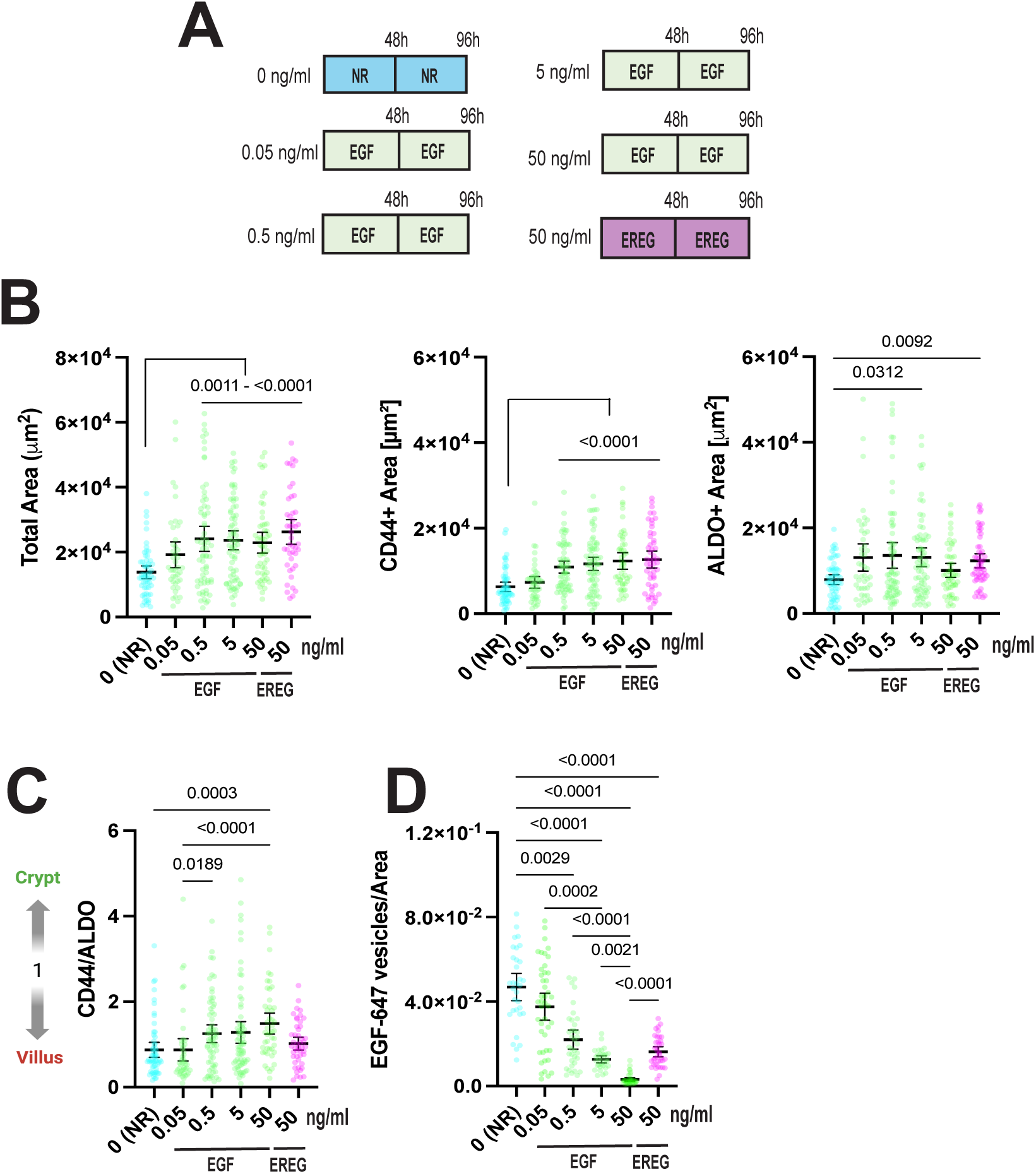
Subsaturating concentrations of EGF trigger both crypt and villus growth. (A-D) Organoids were grown, with NR medium, supplemented with EGF at different concentrations: 50, 5, 0.5 and 0.05 ng/mL or EREG 50 ng/mL for 96h before fixation and immno-staining. (B) Scatter plots showing mean±95%CI, N=3, n=44-71 organoids per condition, significance was calculated by Kruskal-Wallis with Dunn’s multiple comparisons test. p-values <0.05 are shown (C) Relative ratio of CD44 to ALDO areas from (B). Scatter plot mean±95%CI of relative ratio within CD44/ALDO areas. Values above 1 suggests organoid development tends towards crypt proliferation while values below 1 tend to villus differentiation. significance was calculated by Kruskal-Wallis with Dunn’s multiple comparisons test. p-values <0.05 are shown (D) EGF-647 vesicles per area as scatter plots showing mean±95%CI; N=3, n=39-48 crypts/condition; significance was calculated by Kruskal-Wallis with Dunn’s multiple comparisons test. p-values <0.05 are shown

All EGF treatments enhanced organoid total area compared to NR, however crypt growth was significantly enhanced starting at 0.5 ng/mL. Notably, villus differentiation was increased starting at 0.05 ng/mL before being reduced at 50 ng/mL (Figure 3B). Likewise, CD44/ALDO ratio was enhanced towards crypt development starting at 0.5 ng/mL and increased at 50ng/mL as villus area is reduced (Figure 3C).

EGF-647 endocytosis also showed a negative correlation between surface EGFR and EGF concentration. Notably, concentrations that enhanced villus development showed surface EGFR levels similar to EREG treatments (Figure 3D).

### Multiple pulses of EGF deplete EGFR surface expression reducing villus differentiation

Removing saturating amounts of EGF from the culture medium, EREG treatment and sub-saturating EGF concentration all are able to maintain a high level of surface EGFR expression compared to long term EGF treatment. A combination of biosynthesis and endocytic cycling is constantly supplying EGFR to the cell surface, EGF has also been shown to increase EGFR mRNA levels (Scharaw et al., 2016), thus constant turnover is likely critical for the phenotypical outcomes we observe.

To further test this idea, organoids were grown for 96h, however medium was exchanged only 1, 2 or 4 times during this time window in order to constantly deplete surface EGFR in developing organoids (Figure 4A). Total Area of the organoids remained unchanged in EGF and EREG regardless of the number of medium changes, showing that fresh medium addition does not change overall organoid growth (Figure 4B). Contrastingly, Crypt area was increased while Villus area was reduced by multiple EGF medium changes, an effect that was not observed with EREG treatment that maintained Crypt Area in all treatments. 4x EREG treatments significantly reduced Villus Area compared to 96h and 48h, however this change did not significantly affect Crypt/Villus balance (Figure 4C and D). As expected, multiple EGF treatments greatly reduced EGF-647 endocytosis while EREG and NR treatments maintained similar levels of endocytosis in all treatments (Figure 4E).

**Figure 4.**
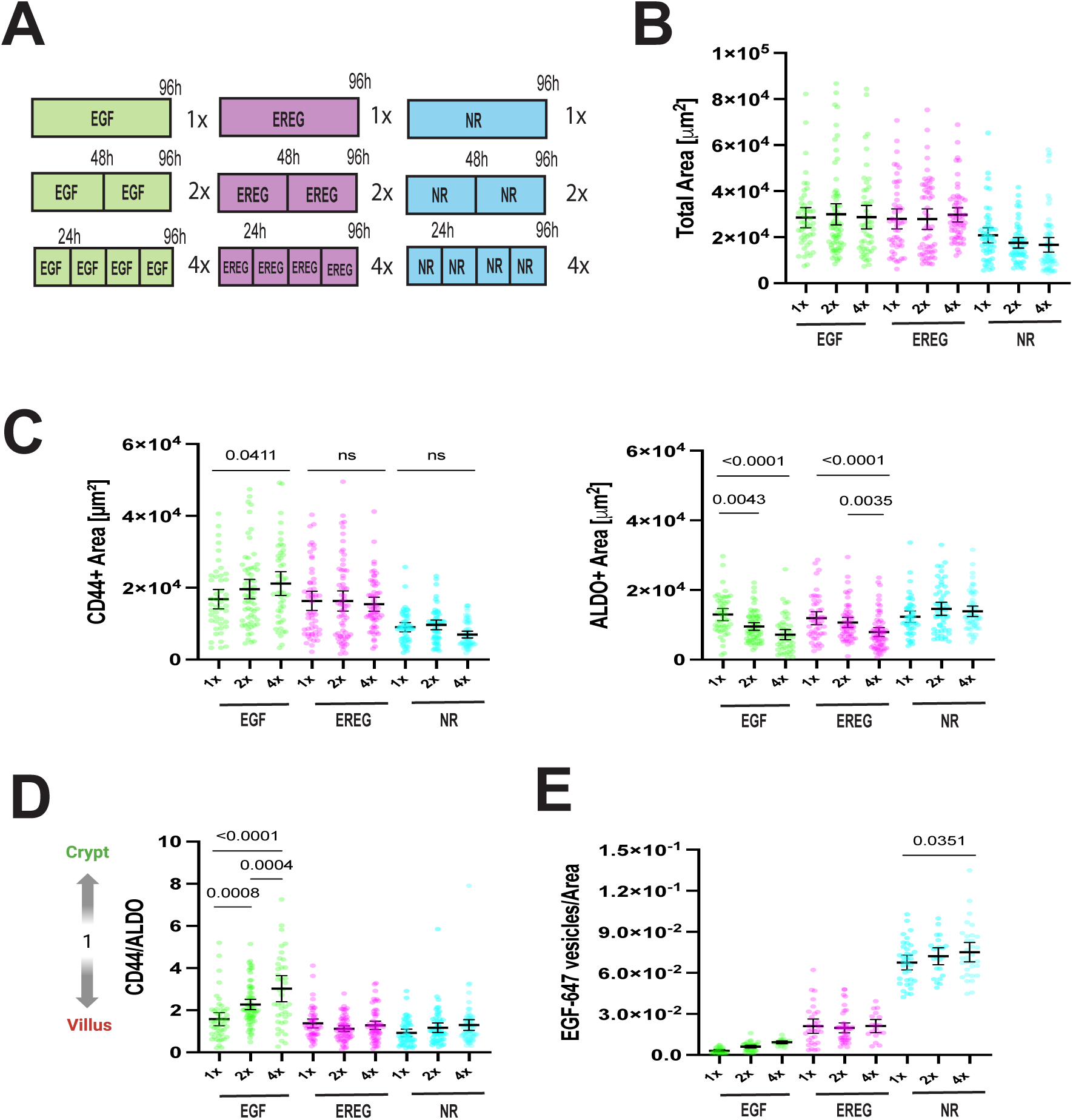
Multiple EGF or EREG medium changes reduces villus differentiation. (A-E) Organoids were grown, with NR medium or supplemented with either EGF (50ng/mL) or EREG (50ng/mL) for 96h with a complete medium change every 24h (4x), 48h (2x) or 96h (1x). (B-C) Scatter plots showing mean±95%CI, N=3, n=47-67 organoids/condition of masked areas corresponding to (B) total Area and (C) CD44+ and ALDO+ areas; significance was calculated by Two-way ANOVA and Sidak’s multiple comparisons test multiple comparisons test. p-values <0.05 are shown. (D) Relative ratio of CD44 to ALDO areas from (C). Scatter plot mean±95%CI of relative ratio within CD44/ALDO areas. Values above 1 suggests organoid development tends towards crypt proliferation while values below 1 tend to villus differentiation. significance was calculated by Two-way ANOVA and Sidak’s multiple comparisons test multiple comparisons test. p-values <0.05 are shown (E) EGF-647 vesicles per area as scatter plots showing mean±95%CI; N=3, n=21-40 crypts/condition; significance was calculated by Two-way ANOVA and Sidak’s multiple comparisons test multiple comparisons test. p-values <0.05 are shown.

### Concluding Remarks

Human plasma levels of EGF range between 0.04 and 0.1 ng/mL (Alshammari et al., 2026), whereas, concentrations of up to 300 ng/mL have been identified in rodent milk and is essential for early gastrointestinal development in newborns. After weaning and with age, EGF levels in the intestine are greatly reduced (Schaudies et al., 1989; Fisher and Lakshmanan, 1990). EGF expression is also upregulated in Paneth cells upon injury in the intestinal epithelia (Calafiore et al., 2023). Absolute concentrations of EGF in tissues could also be obscured by tissue geometry, such as the confined space within an intestinal crypt, allowing for a higher concentration of ligand within a lower volume (Tschumperlin et al., 2004; Schmick and Bastiaens, 2014) In all, the intestinal tissue is exposed to changing of ligands throughout development, with higher levels are often associated with epithelial regeneration processes interspersed within long periods of normal proliferation and differentiation. In turn, periods of low ligand availability enhance long term stability of EGFR expression and enrichment at the plasma membrane, making intestinal crypts susceptible to changes in the concentration of ligands in the environment.

Saturating amounts of EGF might be instrumental for passaging and expanding intestinal organoids, especially in when seeding single stem cells (Boretto et al., 2024). However, seeding single crypts allows for a pre-stablished cellular niche to trigger initial organoid regeneration, even in the absence of medium EGF. Over 48h, the organoids establish a fetal-like morphology which was not influenced by EGF or EREG as organoids developed equally in total area regardless of presence or absence of ligand. Similar developmental profiles could be induced by re-introducing EGF or EREG after 48h and by removing ligands after 1h of incubation upon crypt seeding, highlighting the robustness of EGFR mediated signaling.

To couple phenotype to EGFR activity, we incubated organoids with labelled EGF-647 and examined endocytosis as measurement of the amount of EGFR present in the basolateral membrane after treatments. It is well established in the literature that receptor-bound EGF is released under acidic pH of endosomal compartments. Whether this applies to EREG and EREG-triggered recycling produces ligand-less surface receptors is unknown. Labelled EGF has been extensively used to quantify surface levels and/or total receptor levels in cell lines (Stanoev et al., 2018; Joshi et al., 2023; Roepstorff et al., 2009).This process usually include incubations and washes with ice-cold buffers. These treatments are not possible in organoids where cold washes could compromise the integrity of the Matrigel matrix sustaining organoid morphology and the intracellular cytoskeletal network.

While EGF treatment might promote receptor degradation, thus reducing EGFR endocytosis of EGF-647, it is also possible that smaller vesicles associated to clathrin mediated endocytosis of EGFR monomers could not be detectable by CLSM over the background signal. In other hand, higher levels of surface receptor in NR and EREG treatment enhance non-clathrin mediated endocytosis of EGFR clusters within larger vesicles and higher number of fluorescent molecules which facilitate detection. On the other hand EREG, a low affinity EGFR ligand that has been shown to promote sustained EGFR phosphorylation and differentiation in cultured cells (Freed et al., 2017). After 96h of growth we were unable to observe increased phosphorylation of EGFR in EREG treated organoids, pEGFR levels quantified by immunofluorescence were often comparable to EGF treated organoids, also total levels of EGFR were reduced by EREG 96h treatment. EREG-induced EGFR endocytosis leads to rapid recycling back to the plasma membrane and reduced ubiquitination compared to EGF treatment, however, a fraction of EREG-bound receptors shows colocalization with late endosomal markers (Roepstorff et al., 2009; Deguchi et al., 2024) which could account for degradation.

## Methods

### Mouse intestinal organoids culture

AdDF+++ medium containing Advanced Dulbecco’s modified Eagle’s medium/F12 supplemented with 100 u/mL penicillin/streptomycin, 10 mM HEPES, 2mM Glutamax. ENR medium consists of AdDF+++ supplemented with 1x B27 (Life Technologies) and 1.25 mM N-acetylcysteine, 50ng/mL mEGF (Peprotech, 315-09), 500 ng/mL R-Spondin 1 (Peprotech, 315-32) and 100 ng/mL Noggin (Peprotech, 250-38). Alternatively, medium without EGF was used as “NR medium”. Cryopreserved mouse small intestinal organoids were purchased form STEMCELL Technologies (Cat #70931) and grown in Matrigel 30 μL droplets in 24 well plates in ENR medium for 96-120h before splitting by mechanical disruption with yellow pipette tips. Alternatively for treatments, NR medium was supplemented with 50 ng/mL of EREG (Biotechne, 1195-EP-025,) or 50 ng/mL NRG1 (Biotechne-377-HB-050/CF).

### Immuno-staining

Single organoid crypts were seeded in 20 μL Matrigel droplets in x-well culture chambers 8 wells on coverslips (Sarstedt, 94.6190.802). After 96h of growth organoids were fixed for 40 min in 4% PFA. Then washed 3 times with IF-buffer (0.2% Triton X100, 0.05% Tween20) for 5 min and permeabilized with PBS 0.5% Triton X100 for 1h. Organoids were then blocked with IF-Buffer supplemented with 3% BSA. Alternatively, for staining of phosphorylated EGFR (pEGFR), BSA was replaced by 50% Goat Serum (Invitrogen, 50197Z) and 50% IF buffer for both blocking and antibody staining. Primary antibodies CD44 (1:500; BD Pharmigen™, 560568), Lysozyme (1:1000, DAKO, A0099) and Aldolase (1:500, ab153828, Abcam), pEGFR (1:250,EP774Y,Abcam), Ki-67 (1:500, 55060, BD Biosciences), Rab7 (1:500, D95F2, Cell Signaling Technology) were incubated overnight with agitation at 4°C. Organoids were washed 3 times with IF Buffer for 5 min and then incubated for 1h with Alexa Fluor secondary antibodies (1:500,Thermo-Fisher) and DAPI (1:1000). Alexa Fluor 488 (A11008, A11001), Alexa Fluor 546 (A11081, A11030) and Alexa Fluor 647 (A31573, A21247) secondary antibodies were used. After washing with IF buffer, samples were kept in Clearing Solution (50% v/v Glycerol, 2.5M Fructose) up to imaging.

### Confocal Laser Scanning Microscopy (CLSM) imaging

Organoids were imaged using Leica Confocal TCS SP8 with a 20x/0.75 NA air objective. 2 μm spaced z-stacks were acquired in the fluorescence channels suitable for each sample. Area quantifications were performed using binary masks thresholded from Maximum Intensity Projections of either DAPI, CD44 or AldoB representing total, crypt and villus area respectively. Alternatively, organoid crypts were imaged with 63×1.4 NA oil-immersion objective for pEGFR and CD44 imaging.

### EGF-647 endocytosis

The His-CBD-Intein-(Cys)-hEGF-(Cys) plasmid was kindly provided by Prof. Luc Brunsveld, University of Technology, Eindhoven. EGF-647 was purified in house as previously described (Sonntag et al., 2014; Joshi et al., 2023). For endocytosis in organoids, medium was aspirated from x-well culture chambers, 8 wells on coverslips (94.6190.802, Sarstedt) and rinsed with warm AdDF+++ once. Subsequently, organoids were incubated for 30 min with 50ng/mL of EGF-647 before fixation with 4% PFA. Organoids were imaged in Leica Confocal SP8 with 63x oil immersion objective, crypts were identified by Ki-67 staining. Cellular area was masked with Ki67 and Rab7 staining and the number vesicles were quantified using the find maxima command in Fiji.

### Immunoblotting

Organoid samples were pelleted, resuspended in modified RIPA buffer (50 mM Tris HCl, pH 7.4, 150 mM NaCl, 1 mM EDTA, 1 mM EGTA, 0.5% Triton X-100, and 0.5% sodium deoxycholate) supplemented with Complete Mini EDTA-free protease inhibitor (Roche Applied Science) and phosphatase inhibitor cocktail 2 and 3 (1:100, P5726 and P0044, Sigma–Aldrich), frozen in liquid nitrogen and stored in -70°C freezer. Samples were then dissociated using 10-gauge needle syringes and centrifuged at 12,000 rpm for 15 min at 4°C. Protein concentration was determined by BCA (Thermo Scientific, 23235) and 50 μg of protein was loaded in 8% SDS/PAGE gels and then transferred into PVDF membranes. Membranes were then blocked in intercept buffer and incubated with primary antibody overnight. EGF Receptor (C75B9) (1:1000, Cell Signaling Technology, 2646) for total EGFR, EGFR Y1068 (AP774Y) (1:500; Abcam, 40815) for phospho EGFR (pEGFR), and α-Tubulin (1:10000; Merck, T6074,) were used. Secondary antibodies IRDye 680 donkey anti-Rabbit IgG (1:10000), IRDye 800 donkey anti-Mouse IgG (1:10000) and IRDye 800 donkey anti-Rabbit IgG (1:10000) (LI-COR Biosciences) were used.

### Digital Lightsheet (DLS) Microscopy

Organoids were grown in U-Shaped Capillaries 20 x 1.5 mm (Leica, 158007061) inside Glass-Bottom 35 mm MatTek Dishes (MatTek Life Sciences, P35G-1.5-10-C) for 96h in NR medium. Before imaging, organoids were treated for 1h with 1:500 Cell Mask^TM^ Green Actin Tracking stain (Invitrogen, A57243) and Hoechst 33342 (Invitrogen, C10340 G). Afterwards organoids were washed twice with warm AdDF+++ medium and returned to fresh NR medium. Organoids were imaged in Leica TCS SP8 Digital Lightsheet (DLS) DMi850 using HC PL FLUOTAR 5x/0.15 excitation objective and FLUOTAR L25x/0.95 W DLS detection objective. 50ng/mL of EGF-647 was added after calibration and ROI selection.

### Statistical Analyses

Analysis was performed with at least 3 biological replicates (N=3) unless otherwise stated. Scatter plots show mean±95%CI and bar graphs show mean±SD. For single group comparisons Krustal-Wallis with Dunn’s multiple comparisons test was used. For multiple group comparisons Two-way ANOVA with Sidak’s multiple comparisons test multiple comparisons test. p-values <0.05 are shown. All analyses and graphs were implemented using Prism GraphPad

## Acknowledgments

This project was supported by Max Planck Society for the Advancement of Science. We thank Dr. Malte Schmick for invaluable comments in the production of this manuscript. Special thanks to Dr. Sven Müller for technical support with confocal and DLS microscopy. Figure 1E was created with BioRender.com.

MOC: conceptualization, data curation, formal analysis, experimental methodology, visualization, writing and editing, SS: experimental methodology, editing, LMMN: conceptualization, writing and editing, BS: conceptualization, data curation, visualization, editing, KM: experimental methodology, editing. PHIB: Conceptualization, funding acquisition, supervision, project administration.

Prof. Dr. Philippe I.H. Bastiaens, passed away on May 15, 2025. All authors agree that his inclusion as a co-author is appropriate to honor his intellectual contribution to this work.

The authors state that this research was conducted in the absence of any commercial or financial relationships that could be construed as a potential conflict of interest.

---

**Figure S1.**
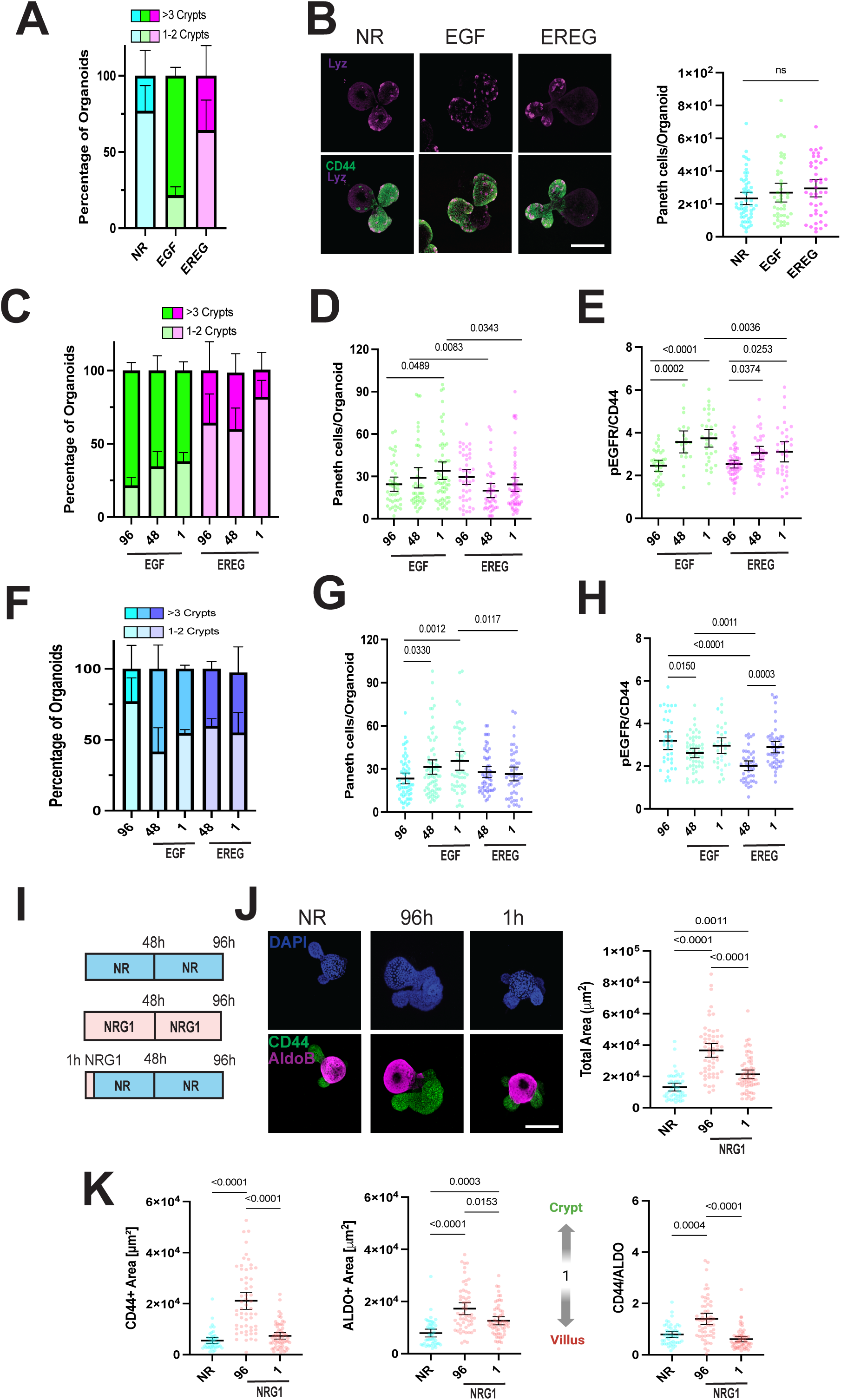
Effect of EGF/EREG temporal modulation in Crypt number, Paneth cell number and pEGFR staining. (A-B) Organoids were grown with NR medium or supplemented with either EGF (50ng/mL) or EREG (50ng/mL) for 96h with a complete medium change at 48h (A) Percentage of Organoids displaying either 1-2 or 3 or more crypts. N=3 (B) (Left) Organoids were immuno-stained with CD44 (Green) and Lysozyme (Lyz, Magenta). Scatter plots showing mean±95%CI of Paneth cells number per organoids was quantified in each condition. Quantification of n=43-56 organoids/condition, N=3, significance was calculated by Krustal-Wallis with Dunn’s multiple comparisons test. p-values <0.05 are shown. (C-E) Organoids were grown, with NR medium supplemented with either EGF (50ng/mL) or EREG (50ng/mL) for 48h or 1h, then complete medium was replaced with NR medium until fixation at 96h. (C) Percentage of Organoids displaying either 1-2 or 3 or more crypts. N=3 (D) Scatter plots showing mean±95%CI of Paneth cells number per organoids was quantified in each condition. Quantification of n=42-61 organoids per condition, N=3, significance was calculated by Two-way ANOVA and Sidak’s multiple comparisons test multiple comparisons test. p-values <0.05 are shown (E) Ratio of pEGFR to CD44 Scatter plots of mean±95%CI. N=3, n=21-49 crypts/condition; significance was calculated by Two-way ANOVA and Sidak’s multiple comparisons test multiple comparisons test. p-values <0.05 are shown. (F-H) Organoids were grown for 1h and 48h with NR, then complete medium was replaced with NR medium supplemented with either EGF (50ng/mL) or EREG (50ng/mL). (F) Percentage of Organoids displaying either 1-2 or 3 or more crypts. N=3 (G) Scatter plots showing mean±95%CI of Paneth cells number per organoids was quantified in each condition. Quantification of n=50-69 organoids/condition, N=3, significance was calculated by Two-way ANOVA and Sidak’s multiple comparisons test multiple comparisons test. p-values <0.05 are shown (H) Ratio of pEGFR to CD44 Scatter plots of mean±95%CI. N=2-3, n=39-54 crypts/condition; significance was calculated by Two-way ANOVA and Sidak’s multiple comparisons test multiple comparisons test. p-values <0.05 are shown. (I) Organoids were grown, with NR medium or NR medium supplemented with NRG1 (50ng/mL) for 96h or 1h. After 1h complete medium was replaced with NR medium until fixation at 96h. (J) Organoids were immuno-stained with CD44 (Green) AldoB (Magenta) and DAPI (Blue). Scale Bar 100 μm. (J-K) Scatter plots showing mean±95%CI of masked areas corresponding to total, CD44+, ALDO+ and ratio Quantification of n=58-65 organoids/condition, N=2-3, significance was calculated by Krustal-Wallis with Dunn’s multiple comparisons test. p-values <0.05 are shown.

**Figure S2.**
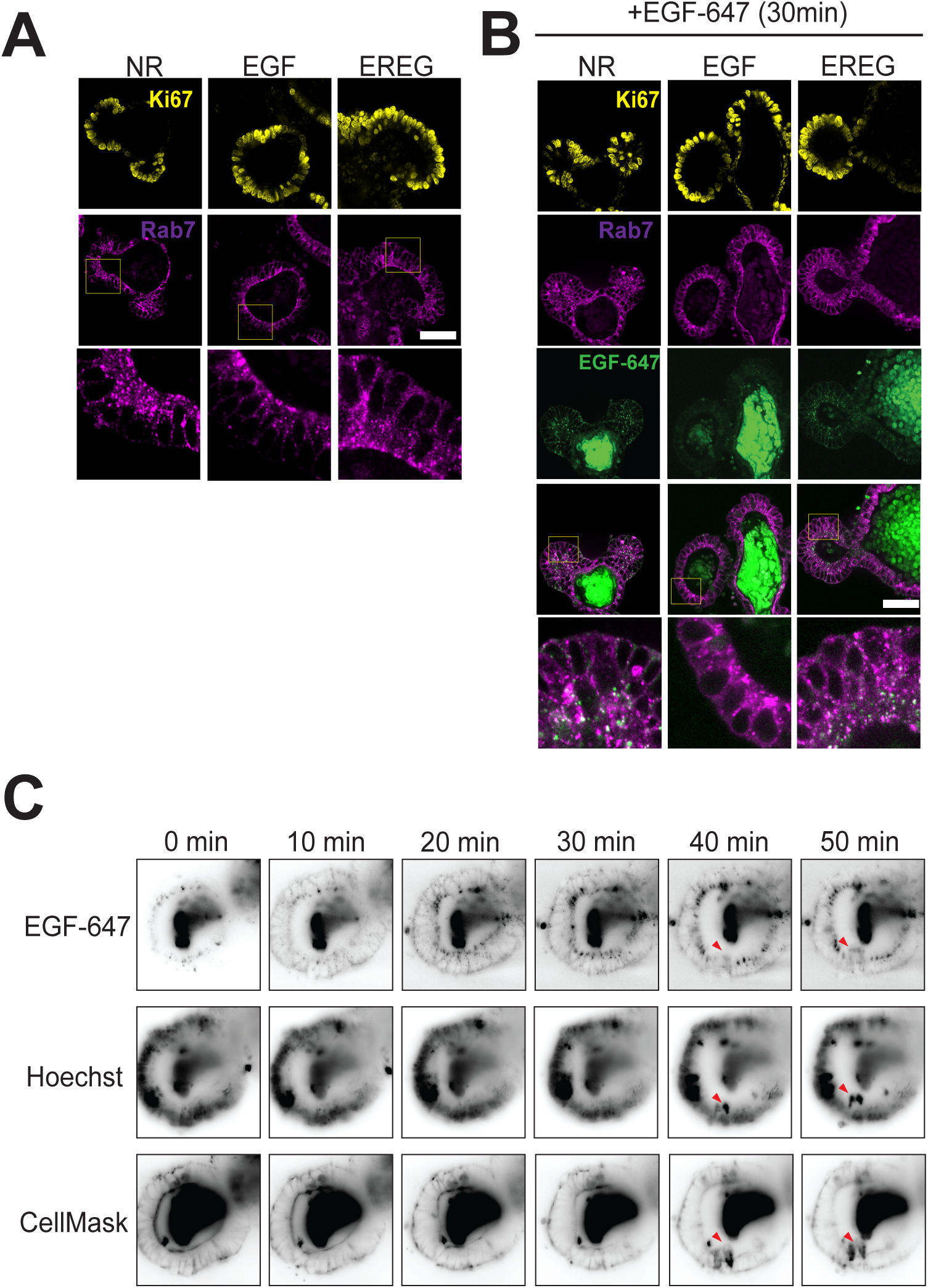
Organoids grown for 48h often display no villus region and retain EGF-647 endocytosis in proliferating cells. (A) Brightfield time lapse imaging of developing organoid from seeded crypts in EGF medium showing standard developmental trajectories between seeding (0h), 24h, 48h, and 96h (B) Organoids were grown, with NR medium or supplemented with either EGF (50ng/mL) or EREG (50ng/mL) for 48h before fixing and staining. (C) Percentage of Organoids showing ALDO+ villus region after 48h of development, N=3. (D) Scatter plots of masked areas corresponding to total area, for all organoids (Left) and organoids with ALDO+ region showing mean±95%CI, N=3, n=45-70 organoids/condition, (Right). Quantification of, significance was calculated by Kruskal-Wallis with Dunn’s multiple comparisons test. p-values <0.05 are shown, ns non-significant. (E) Organoids were immuno-stained with CD44 (Green) AldoB (Magenta) and DAPI (Blue). Organoids ALDO- and ALDO+ are shown for each treatment. Scale Bar 100 μm. (F) Alexa Fluor-647 labelled EGF (50ng/mL) is added to organoids after 48h of growth, after 30min of incubation organoids were fixed and stained with Ki67 (Yellow), Rab7 (Magenta) and DAPI (Blue). EGF-647 is show in (Green). Representative images of Round or Crypt morphologies are represented for each treatment.

**Figure S3.**
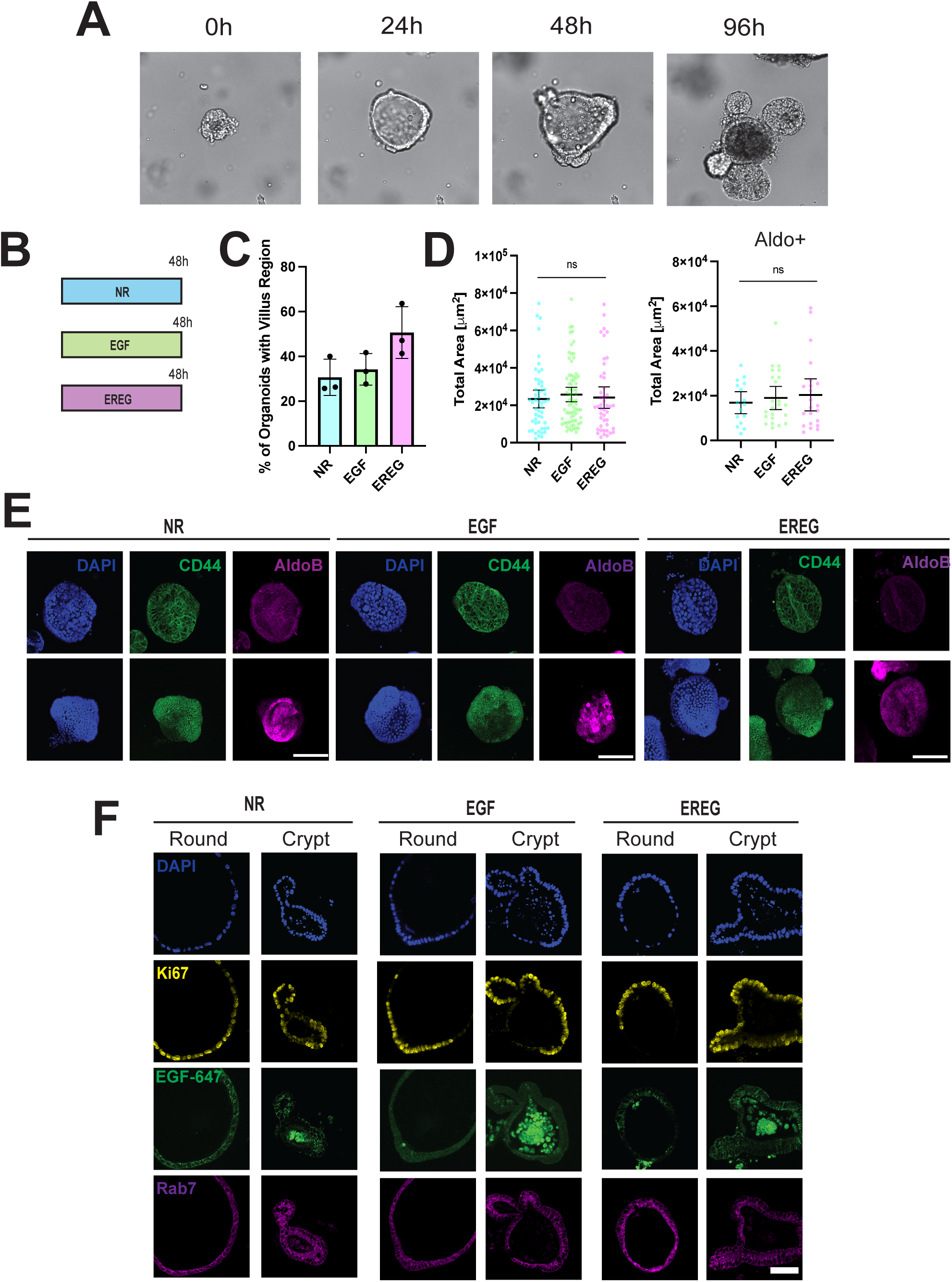
EGF-647 endocytosis in organoid crypts co-localizes with late endosomal marker Rab7. (A-B) Organoids were grown, with NR medium or supplemented with either EGF (50ng/mL) or EREG (50ng/mL) for 96h with a complete medium change at 48h. (A) Organoids were immuno-stained for Ki67 (Yellow) and Rab7 (Magenta), Yellow boxes denote zoom-in areas shown in the row below. (B) 50ng/mL of Alexa Fluor-647 labelled EGF (Green) is added to organoids after 96h of growth, after 30min of incubation organoids were fixed and stained. Yellow boxes denote zoom-in areas shown in the row bellow with EGF-647 (Green) and Rab7 (Magenta). (C) Digital Lightsheet Microscopy was used to see EGF-647 endocytosis in an organoid crypt grown for 96h in NR medium. Images showing Cell Mask, Hoechst and EGF-647 every 10 minutes. Red triangles show cells protruding into the lumen. For more details see *Methods* and (Video1)

Video1. EGF-647 endocytosis in NR grown organoid crypt using Digital Light Sheet imaging.

